# Evolclust: automated inference of evolutionary conserved gene clusters in eukaryotes

**DOI:** 10.1101/698621

**Authors:** Marina Marcet-Houben, Toni Gabaldón

## Abstract

**Motivation:** The evolution and role of gene clusters in eukaryotes is poorly understood. Currently, most studies and computational prediction programs limit their focus to specific types of clusters, such as those involved in secondary metabolism.

**Results:** We present Evolclust, a python-based tool for the inference of evolutionary conserved gene clusters from genome comparisons, independently of the function or gene composition of the cluster. Evolclust predicts conserved gene clusters from pairwise genome comparisons and infers families of related clusters from multiple (all vs all) genome comparisons.

**Availability:** https://github.com/Gabaldonlab/EvolClust/

## 1 Introduction

Gene order in eukaryotic genomes tends to be poorly conserved through evolution (Dávila López et al., 2010). Despite this trend, certain groups of genes remain close in the genome over long evolutionary distances, which suggests that selection acts to maintain their genomic co-localization. Genes within such conserved clusters may have functional links (Lee and Sonnhammer, 2003; Wisecaver et al., 2014), and can be transcribed in a coordinated manner (Boutanaev et al., 2002; Reimegård et al., 2017). Comparative genomics can be used to uncover groups of genes that remain significantly closer than expected, despite extensive gene order shuffling. Using this rationale, we developed Evolclust, an algorithm that detects groups of neighboring genes shared by compared genomes. Unlike most currently available gene cluster prediction algorithms, Evolclust is able to deal with eukaryotic complexity (i.e. multiple chromosomes and extensive duplication patterns), is prepared to perform large scale comparisons using hundreds of genomes, and is not limited to a specific type of gene clusters, such as secondary metabolite clusters (Khaldi et al., 2010; Medema et al., 2011).

## 2 Algorithm

### 2.1 Cluster definition

In Evolclust a conserved cluster is defined as a group of neighboring genes whose homologous genes are also neighboring each other in a different genome. Genes with homologs in the clusters defined in the two genomes are deemed cluster genes. Cluster genes do not need to be strictly in the same order in the two genomes considered, and up to three non-cluster genes (i.e. without homologs in the corresponding cluster defined in the other genome) are allowed in between any two cluster genes. The cluster needs to contain genes encoding at least four different protein families so as to avoid identifying clusters composed solely of tandemly repeated genes.

### 2.2 Evolclust pipeline

Evolclust is designed to run parts of the pipeline in a computing cluster, enabling the processing of a large number of genomes. For smaller databases it also provides an option to perform all steps with a single command. As an output Evolclust provides lists of inferred clusters grouped into families. One of the inherent drawbacks of Evolclust is that it detects conserved clusters by comparison against background gene order conservation. Therefore it will not always be able to predict clusters found exclusively in closely related species that have overall conserved gene order throughout the entire genome.

Evolclust algorithm proceeds as follows (see documentation for further details: https://github.com/Gabaldonlab/EvolClust/wiki). The starting file is a list of families of homologous proteins. Based on pairs of homologous proteins all possible clusters between a pair of species is calculated. The algorithm used to calculate clusters can be found in figure 3 of the documentation. A conservation score (C) is calculated for all found conserved regions as follows: C=[(m^2^+n^2^)−(o^2^−p^2^)]/2(m+n+o+p); where m and n are the number of shared genes for each region (i.e. those with homologs in the other region), and o and p the genes specific to each region. C is calculated across the whole conserved region and iteratively for smaller subregions so as to account for all possible cluster sizes. The distribution of C values in each pairwise comparison provides a measure of the expected conservation of randomly chosen genes between two species.

Once the threshold values are calculated the algorithm iterates through the predicted regions and selects all possible clusters with C above the threshold. Clusters are predicted for each pair of species. Hence, some redundancy is expected. Such redundancy is lowered by collapsing significantly overlapping gene clusters predicted in a given species. Once all clusters are defined for all species, C values between all of them are calculated as explained above. These scores are then used to group clusters into families using the mcl algorithm (Enright et al., 2002). A subsequent cleaning step trims clusters based on the average length of the family. Finally, the initial list of clusters is sourced again and homologous clusters are added to a family if they share, with any cluster of that family, 90% of homologous proteins and have a C>0.95 indicating they were likely discarded because they were in closely related species.

## 3 Benchmarking

To assess whether Evolclust is able to recall meaningful gene clusters we obtained a set of known secondary metabolism gene clusters that are found in at least two fungal genomes (see supplementary table 1). The final list contained 26 experimentally characterized cluster families that comprised a total of 97 individual gene clusters.

Evolclust was first run among the genomes of 341 fungal genomes. A total set of 12,120 cluster families that contained 118,699 clusters was predicted. Out of the 97 clusters 88 (91% of the total set) were found, though for some of them the exact boundaries of the cluster were not detected. In total, 31 clusters (32%) were predicted with exact boundaries, while 57 (59%) had discrepancies with the defined benchmark cluster. However boundary differences were generally low. The median number of extra genes was of 1, while the median number of missing genes was 0.

We compared our results with those obtained in the same set of genomes when using the popular SMURF (Khaldi et al., 2010) and ANTISMASH (Medema et al., 2011) algorithms, which are designed to detect this specific kind of gene clusters (see table S2 and S3 respectively). SMURF was able to find 79 (81%) of the clusters and ANTISMASH found 86 (88%). SMURF predicted 8 correct clusters and had a median of 8 additional genes, ANTISMASH had a single correctly predicted cluster and a median of 11 additional genes. Thus Evolclust is able to detect the presence of most (91%) known conserved clusters

We compared Gecko3 (Winter et al., 2016) and Evolclust in a smaller, randomly chosen subset of 30 genomes (see supplementary material). We observed that both methods predicted a similar number of gene clusters, and had a 77% of overlap in terms of predicted clusters. In addition 87% of the clusters predicted by both methods were assigned to the same families. A final test was made running EvolClust on randomly shuffled genomes. We found that only 1 of the 6482 predicted clusters appeared in only one of the randomly shuffled genomes. Collectively, these results show that Evolclust provides an alternative method to calculate conserved gene clusters that are biologically relevant and is able to process massive amounts of data.

