## Supplementary for "Evolclust: automated inference of evolutionary conserved gene clusters in eukaryotes"

**Supplementary text: 1**

1.- Comparison between Evolclust and Gecko3. 1

1.1.- Introduction 1

1.2.- Cluster Comparison 2

1.3.- Computation time and resources. 4

2.- Comparison to random genomes. 4

**Supplementary figures 5**

**Supplementary tables 8**

**References: 23**

### Supplementary text:

#### 1.- Comparison between Evolclust and Gecko3.

##### 1.1.- Algorithmic differences

Gecko3 (v3.1) (Winter *et al.*, 2016) is an automated method to detect conserved clusters in a given set of genomes. It allows the user to detect clusters using only one genome as reference or do i based on an all by all comparison. Even when using the all by all comparison, there is a cluster in the family, which is designated as the reference. Clusters are grouped into families based on a calculated distance to this reference cluster. This distance is calculated by computing the number of changes a cluster needs to undertake to have the same gene content as the reference cluster. Clusters are finally filtered based on a p-value calculated against a randomly shuffled genome. The user is able to provide different parameters which will regulate the distance (-d), the number of times a cluster appear in order to be considered (-q) and the minimum length of the reference cluster (-s). It is of note that the size requirement does not apply to all members of the cluster, so even if the size has been set to 5, there may be members of the family of smaller sizes.

From a theoretical standpoint Evolclust and Gecko3 differ in mainly two ways: i) the use of one of the clusters as reference even in an all by all analysis, and ii) the way in which clusters are distinguished from non-significant gene order conservation.

The use of a reference can directly affect the results of the cluster families. Take the cluster shown in Figure S1A and assume we allow a maximum distance of 3. In the first panel Species 1A is chosen as the reference cluster, and the clusters in species 1A and species 2A will span the entirety of the cluster depicted in the image. For species 3A, the cluster will stop at the presence of the second non-homologous protein, shown in the image as a white square, due to the fact that the distance becomes higher than 3. In the second panel, the reference cluster is the one in species 3A, in this case, proteins F, G and H are never included in the cluster since their inclusion when comparing to the reference will result in both cases in distances above 3. As Gecko3, Evolclust also has limitations in how it computes boundaries, but instead of choosing a cluster for reference it will calculate the representation of the individual proteins in a family. Proteins represented in less than ⅓ of the family will be discarded.

The second difference regards the statistics that help distinguish clusters from non-significant gene order conservation. Gecko3 simply calculates a p-value against a randomly shuffled genome that defines whether the group of genes included in the reference cluster could be observed together by chance. Clusters also need to be present in a certain number of genomes in order to be considered (quorum parameter). Still, it does not specifically search to distinguish between a potential functional cluster and what is considered an expected level of gene order conservation, which would vary according to the evolutionary distance of the compared genomes. In contrast, Evolclust calculates the actual background level of gene order conservation between pairs of genomes and uses this information to discriminate between clusters and conserved gene order. Given two related genomes with high levels of conserved gene order, Evolclust will not be able to provide a list of clusters whereas Gecko will provide a list, although their significance would be questionable because two closely related genomes are expected to have a higher gene order conservation than two randomly shuffled genomes.

##### 1.2.- Comparison of predicted clusters

We attempted to use Gecko3 in the prediction of secondary metabolism gene clusters as we had done with Evolclust, but after 21 days running on a 32Gb of RAM machine we still obtained no results (-q 2 -s 3 -d 3) therefore we decided to selected a set of 30 fungal genomes from the full dataset. The list of genomes and their taxonomy can be seen in table S4.

We ran Evolclust and Gecko3 on this set of genomes. For Evolclust we used the following parameters: minimum cluster size 5, maximum cluster size 35, threshold method of two standard deviations, and 3 contiguous non-homologous proteins. Gecko3 was run several times with different quorum and distance parameters: gecko-q3d3 (-s 5 -d 3 -q 3), gecko-q4d3 (-s 5 -d 3 -q 4), gecko-q5d3 (-s 5 -d 3 -q 5) and gecko-q4d5 (-s 5 -d 5 -q 4). Only clusters which passed their filters were considered. Cluster families were also filtered to delete clusters with a size below 5. If a family has only one cluster remaining after this filter, the family was deleted. This last filter, resulted in a reduction of the predicted cluster families by Gecko3 that ranged between 49% and 54% of the families.

The number of predicted families between the two methods is comparable. Evolclust detects 1691 families with a total of 6482 individual clusters. Gecko3 predictions obtain between 1308 and 2107 families (gecko-q5d3 and gecko-q3d3) and between 5528 and 7468 individual clusters. There is though a notable difference in cluster sizes. As seen in Figure S2, Gecko rarely predicts clusters with a size above 20 (roughly between 0.5% and 1.6% of the clusters are larger) whereas in Evolclust 9% of the clusters are above this size.

We now compare the clusters themselves, to see whether the two methods predict the same clusters or if there are differences. For this, we compared all clusters individually and calculated the overlap between them. The clusters are divided in different categories depending on the level of overlap: identical: all proteins are shared between two clusters, 100% one cluster is a subset of the other one, and 90%, 75%, 50% and 30% where at least one of the clusters shares a given percentage with the other cluster. To ensure that the comparison works, we first compared two different runs of Gecko3 (gecko-q3d3 versus gecko-q5d3). As expected, we find that a large percentage of clusters are shared between the two runs: 98% of the clusters in gecko-q5d3 are found in gecko-q3d3, and 73% of clusters in gecko-q3d3 are found in gecko-q5d5. These results are congruent with the difference made by the quorum parameter and the fact that there are 1940 clusters more predicted in gecko-q3d3. Also as expected, a large percentage of the shared clusters are identical between the two runs of Gecko3 (80.4%) and when not identical in many cases one cluster is a subset of the other one (13.0%).

We now compare the results obtained in Evolclust to Gecko3, we will comment on the gecko-q4d3 run as it is the one with a similar number of clusters (6482 clusters in Evolclust versus 6603 clusters in gecko-q4d3), still, all comparisons can be found in table S5. Considering all levels of overlap, we observe that Evolclust and gecko-q4d3 share approximately 76% of the clusters, of which only ~12% are identical. Still, a large percentage of clusters are found in the 100% category where one cluster is the subset of another one (48%). This is congruent with our previous observation that clusters predicted by Gecko tend to be smaller.

Another interesting measure is whether the clusters, when predicted by both methods, are grouped into families in the same way. In the three comparisons we observed that 87% of the families predicted by one method correspond to a single family in the other method, 10% of the remaining families appear split in two. So, we can conclude that, when clusters are predicted by both methods, they tend to group in the same families.

Differences observed in cluster prediction can likely be attributed to the differences in the prediction method. In Figure S3 we show an example of overlapping clusters predicted by Evolclust (Figure S3a) and two clusters in gecko-q4d3 (Figure S3 b and c). As seen when comparing the two predictions, neither of the methods is perfect. Gecko3 splits the cluster in two even though the genes that are found in between are also conserved in most species and an argument could be made on the conservation of this region as a single cluster. Yet the first family (Figure S3b) includes a pair of genes (8850 and 2918) that have been excluded from Evolclust.

##### 1.3.- Computation time and resources.

For this dataset, with only one thread, Evolclust took 32 hours to complete. This process could be sped up when using a computation cluster as part of the processes can be parallelized. This would result is a faster run yet it would require a higher degree of user interaction. Still it opens the possibility to process much larger datasets in a more efficient way.

The time required to run Gecko3 was dependent on the parameters, so for the runs of gecko-q3d3, gecko-q4d3 and gecko-q5d3 only 2 to 3 hours were needed. An increase of the d parameter to 5 already resulted in an increase in computation time to 7 - 9 hours. So, while Gecko3 is more efficient than Evolclust, in this dataset, the fact that Evolclust can be parallelized will provide a clear advantage in computation time in increasingly large datasets.

##### 2.- Comparison to randomly shuffled genomes.

One possible drawback of the method is that clusters including multiple proteins from large protein families are included at random. In order to ensure that this was not the case, we proceed as follows: using the evolclust predictions obtained for the comparison with Gecko3, for each genome included, we build 100 randomly shuffled genomes and then we searched whether the clusters found in these genomes were also found in the randomized genomes. Out of 6482 clusters found in the dataset, only 1 cluster was predicted in one of the random genomes. This shows, that although the possibility for random clusters exists, it is very limited and as such not considered. The script needed to double check for spurious predictions is included in the github repository under the additional scripts folder (https://github.com/Gabaldonlab/EvolClust/blob/master/additional_scripts/compare_to_random_genomes.py).

#### Supplementary figures

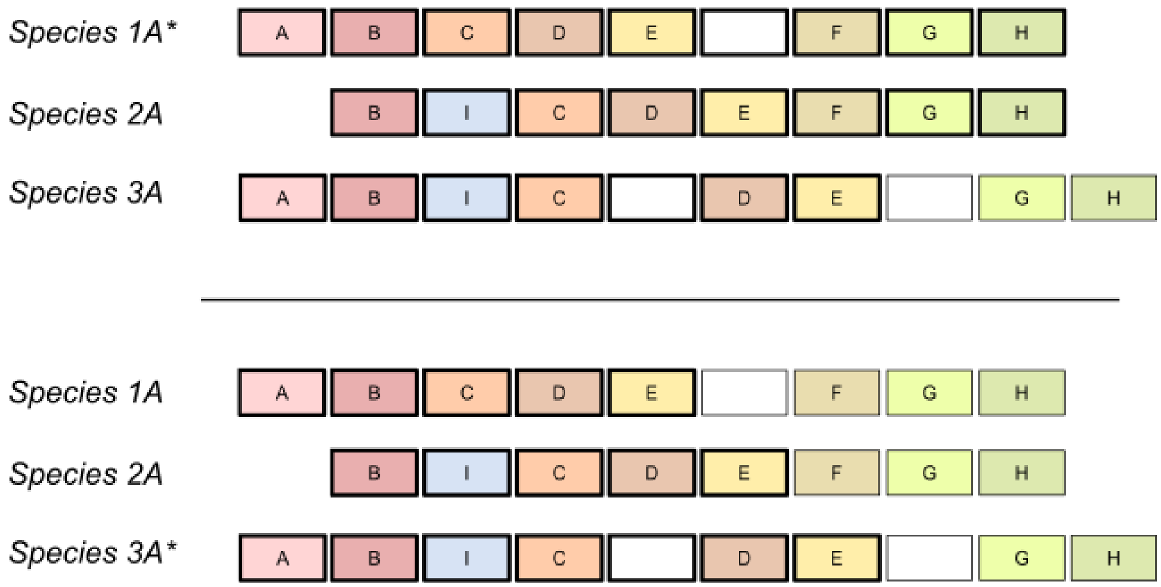


Figure S1: Difference in cluster prediction with Gecko3 depending on the chosen reference. Each gene is represented by a colored square, same colours and numbers indicate the same protein family. Each line belongs to a different species which is specified to the left of the cluster.

Squares with a thick black border indicate the genes that will be part of the predicted cluster. White squares indicate non-homologous proteins.

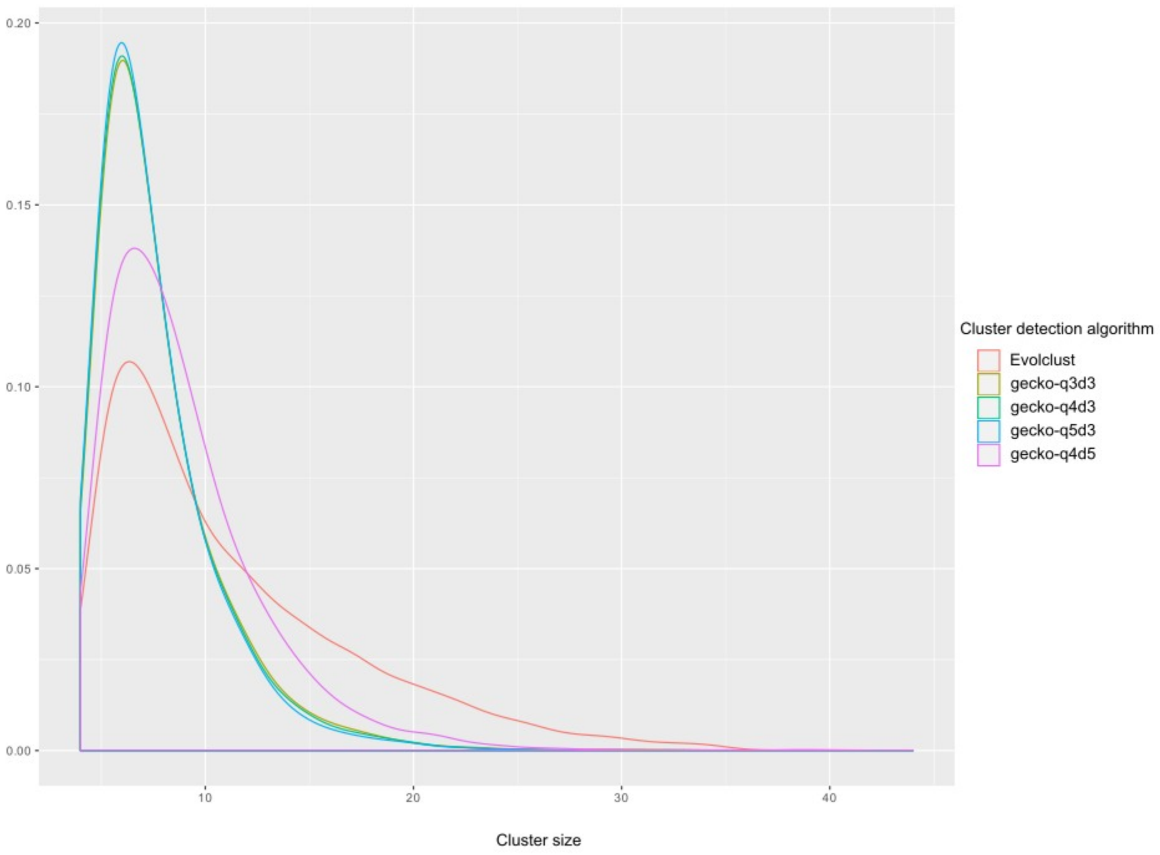


**Figure S2: Distribution of cluster sizes as predicted by Gecko and Evolclust.**

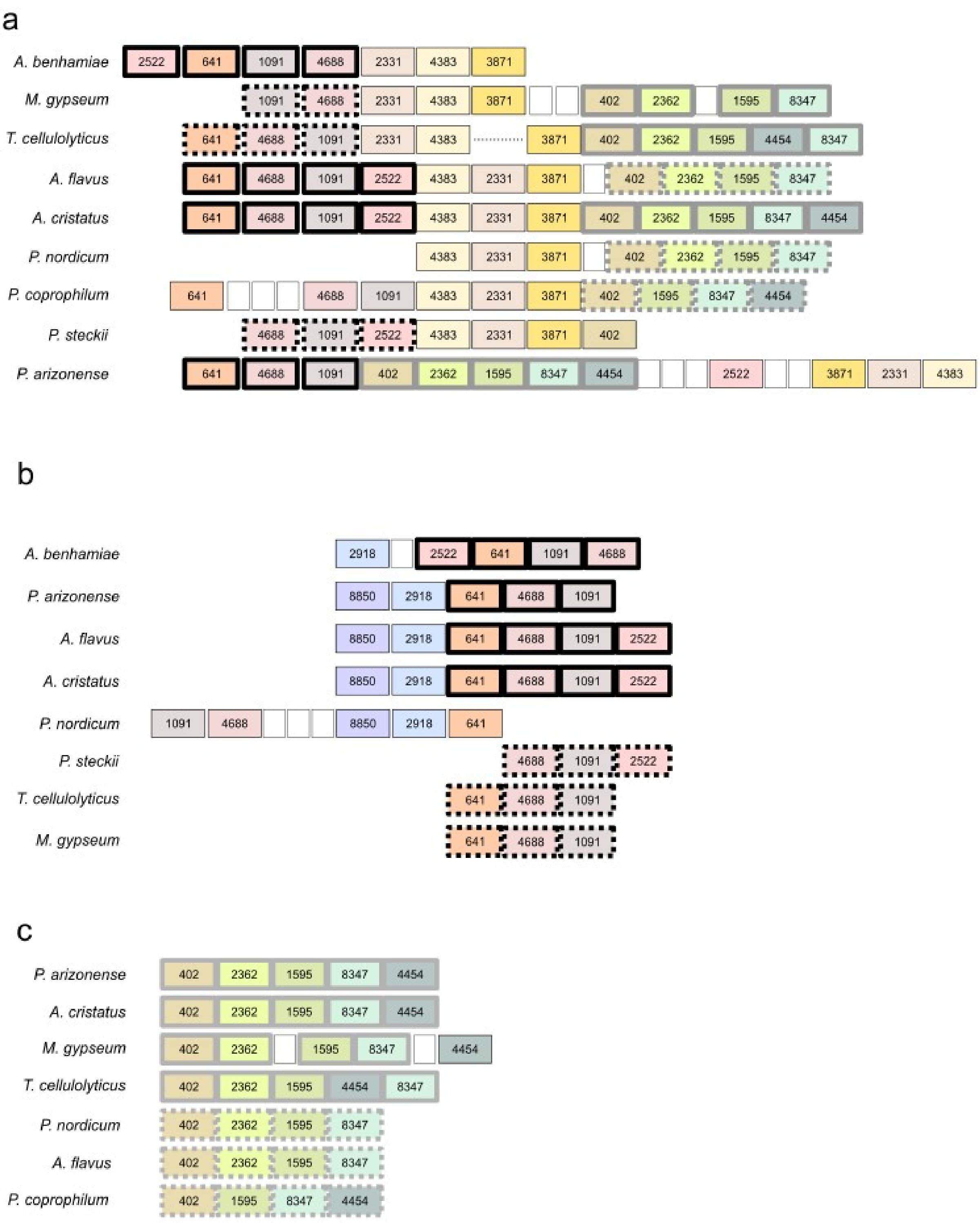


**Figure S3: Comparison of a cluster predicted with Evolclust and two overlapping clusters predicted by Gecko3**

a.- Cluster predicted by Evolclust. Each gene is represented by a colored square, same colours and numbers indicate the same protein family. Each line belongs to a different species which is specified to the left of the cluster. Squares with a thick black border are shared with the first cluster family predicted by gecko-q4d3. If the border is dotted the cluster predicted in gecko-q4d3 does not fulfill the minimum size. Similarly, squares surrounded by a grey border match proteins found in the second gecko-q4d3 cluster family. Smaller white squares indicate non-homologous proteins. b.- First cluster detected by gecko-q4d3. c.- Second cluster detected by gecko-q4d3.

### Supplementary tables

**Table S1: Prediction of secondary metabolism gene clusters with Evolclust.**
Columns indicate the name of the cluster, the pubmed ID of the paper describing the cluster, the species in which it is predicted, a tag indicating whether the cluster was predicted with the correct boundaries (Perfect), whether it was partially predicted (Partial) or not predicted. When predicted, information is provided about the predicted cluster size, the number of additional genes and the number of missing genes.

| Cluster name | PUBMED ID | Species name | Evolclust | | | |
|---|---|---|---|---|---|---|
| | | | **TAG** | **Size** | **Additional genes** | **Missing genes** |
| asperfuranone | 19199437 | *Aspergillus terreus* | Perfect | 8 | 0 | 0 |
| asperfuranone | 19199437 | *Penicillium antarcticum* | Perfect | 8 | 0 | 0 |
| aspernidineA | 23706169 | *Aspergillus calidoustus* | Partial | 6 | 2 | 0 |
| aspernidineA | 23706169 | *Emericella nidulans* | Perfect | 6 | 0 | 0 |
| aurofusarin | 16879655 | *Fusarium poae* | Partial | 8 | 0 | 4 |
| aurofusarin | 16879655 | *Fusarium pseudograminearum* | Partial | 8 | 0 | 4 |
| bikaverin | 23308280 | *Fusarium fujikuroi* | Partial | 6 | 1 | 0 |
| bikaverin | 23308280 | *Fusarium mangiferae* | Partial | 6 | 1 | 0 |
| bikaverin | 23308280 | *Fusarium oxysporum* | Partial | 6 | 1 | 0 |
| bikaverin | 23308280 | *Fusarium verticillioides* | Partial | 6 | 1 | 0 |
| butenolide | 17175185 | *Fusarium avenaceum* | Partial | 8 | 1 | 0 |
| butenolide | 17175185 | *Fusarium graminearum* | Partial | 8 | 2 | 0 |
| butenolide | 17175185 | *Fusarium poae* | Partial | 8 | 3 | 0 |

| | | | | | | |
|---|---|---|---|---|---|---|
| butenolide | 17175185 | *Fusarium pseudograminearum* | Partial | 8 | 3 | 0 |
| depudecin | 19737099 | *Colletotrichum orchidophilum* | Perfect | 6 | 0 | 0 |
| depudecin | 19737099 | *Microsporum gypseum* | Perfect | 6 | 0 | 0 |
| FDB2 | 26808652 | *Fusarium fujikuroi* | Partial | 11 | 0 | 5 |
| FDB2 | 26808652 | *Fusarium mangiferae* | Partial | 11 | 0 | 5 |
| FDB2 | 26808652 | *Fusarium proliferatum* | Partial | 11 | 0 | 5 |
| fumitremorgin | 16755625 | *Aspergillus fumigatus* | Partial | 7 | 0 | 2 |
| fumitremorgin | 16755625 | *Neosartorya fischeri* | Partial | 7 | 0 | 2 |
| fusaric_acid | 22652150 | *Fusarium fujikuroi* | Not_found | | | |
| fusaric_acid | 22652150 | *Fusarium mangiferae* | Not_found | | | |
| fusaric_acid | 22652150 | *Fusarium proliferatum* | Not_found | | | |
| fusarin | 22652150 | *Fusarium fujikuroi* | Perfect | 9 | 0 | 0 |
| fusarin | 22652150 | *Fusarium proliferatum* | Perfect | 9 | 0 | 0 |
| fusarubin | 22492438 | *Fusarium avenaceum* | Partial | 5 | 0 | 1 |
| fusarubin | 22492438 | *Fusarium fujikuroi* | Partial | 5 | 0 | 1 |
| fusarubin | 22492438 | *Fusarium mangiferae* | Partial | 5 | 0 | 1 |
| fusarubin | 22492438 | *Fusarium oxysporum* | Partial | 5 | 0 | 1 |
| fusarubin | 22492438 | *Fusarium proliferatum* | Partial | 5 | 0 | 1 |
| gibberellin | 9917370 | *Fusarium fujikuroi* | Not_found | | | |
| gibberellin | 9917370 | *Fusarium mangiferae* | Not_found | | | |
| gibberellin | 9917370 | *Fusarium proliferatum* | Not_found | | | |
| gliotoxin | 15979823 | *Aspergillus fumigatus* | Partial | 10 | 0 | 2 |
| gliotoxin | 15979823 | *Aspergillus udagawae* | Partial | 11 | 1 | 1 |
| gliotoxin | 15979823 | *Neosartorya fischeri* | Partial | 11 | 0 | 1 |
| gliotoxin | 15979823 | *Neosartorya udagawae* | Partial | 11 | 1 | 1 |

| | | | | | | |
|---|---|---|---|---|---|---|
| HAS | 25386169 | *Arthroderma benhamiae* | Partial | 7 | 0 | 1 |
| HAS | 25386169 | *Arthroderma otae* | Partial | 7 | 0 | 1 |
| HAS | 25386169 | *Aspergillus fumigatus* | Perfect | 8 | 0 | 0 |
| HAS | 25386169 | *Aspergillus udagawae* | Perfect | 8 | 0 | 0 |
| HAS | 25386169 | *Neosartorya fischeri* | Perfect | 8 | 0 | 0 |
| HAS | 25386169 | *Neosartorya udagawae* | Perfect | 8 | 0 | 0 |
| HAS | 25386169 | *Trichophyton equinum* | Partial | 7 | 0 | 1 |
| HAS | 25386169 | *Trichophyton soudanense* | Partial | 7 | 0 | 1 |
| Helvonic acid | 19415934 | *Aspergillus fumigatus* | Partial | 8 | 0 | 1 |
| Helvonic acid | 19415934 | *Aspergillus udagawae* | Partial | 7 | 0 | 2 |
| Helvonic acid | 19415934 | *Metarhizium anisopliae* | Partial | 8 | 0 | 1 |
| Helvonic acid | 19415934 | *Metarhizium brunneum* | Perfect | 9 | 0 | 0 |
| Helvonic acid | 19415934 | *Metarhizium guizhouense* | Perfect | 9 | 0 | 0 |
| Helvonic acid | 19415934 | *Metarhizium robertsii* | Perfect | 9 | 0 | 0 |
| Helvonic acid | 19415934 | *Neosartorya fischeri* | Partial | 8 | 0 | 1 |
| Neosartoricin | 23758576 | *Arthroderma benhamiae* | Perfect | 5 | 0 | 0 |
| Neosartoricin | 23758576 | *Arthroderma otae* | Perfect | 5 | 0 | 0 |
| Neosartoricin | 23758576 | *Aspergillus fumigatus* | Perfect | 5 | 0 | 0 |
| Neosartoricin | 23758576 | *Microsporum gypseum* | Perfect | 5 | 0 | 0 |
| Neosartoricin | 23758576 | *Neosartorya fischeri* | Perfect | 5 | 0 | 0 |
| Neosartoricin | 23758576 | *Trichophyton rubrum* | Perfect | 5 | 0 | 0 |
| Neosartoricin | 23758576 | *Trichophyton soudanense* | Perfect | 5 | 0 | 0 |
| Neosartoricin | 23758576 | *Trichophyton tonsurans* | Perfect | 5 | 0 | 0 |
| Neosartoricin | 23758576 | *Trichophyton verrucosum* | Perfect | 5 | 0 | 0 |
| Neosartoricin | 23758576 | *Trichophyton violaceum* | Perfect | 5 | 0 | 0 |

| | | | | | | |
|---|---|---|---|---|---|---|
| NG391 | 27801295 | *Metarhizium anisopliae* | Partial | 6 | 3 | 0 |
| NG391 | 27801295 | *Metarhizium brunneum* | Partial | 6 | 3 | 0 |
| NG391 | 27801295 | *Metarhizium guizhouense* | Partial | 6 | 3 | 0 |
| NG391 | 27801295 | *Metarhizium majus* | Partial | 6 | 3 | 0 |
| NG391 | 27801295 | *Metarhizium robertsii* | Partial | 6 | 3 | 0 |
| Patulin | 19383676 | *Aspergillus clavatus* | Partial | 14 | 0 | 1 |
| Patulin | 19383676 | *Penicillium expansum* | Perfect | 15 | 0 | 0 |
| Perithecial pigment | 22652150 | *Fusarium avenaceum* | Partial | 4 | 1 | 1 |
| Perithecial pigment | 22652150 | *Fusarium fujikuroi* | Partial | 4 | 1 | 1 |
| Perithecial pigment | 22652150 | *Fusarium mangiferae* | Partial | 4 | 1 | 1 |
| Perithecial pigment | 22652150 | *Fusarium oxysporum* | Partial | 4 | 1 | 1 |
| Perithecial pigment | 22652150 | *Fusarium proliferatum* | Partial | 4 | 1 | 1 |
| pigments | 10515939 | *Aspergillus clavatus* | Not_found | | | |
| pigments | 10515939 | *Neosartorya fischeri* | Not_found | | | |
| PKI | 23617571 | *Aspergillus rambellii* | Partial | 7 | 0 | 1 |
| PKI | 23617571 | *Aspergillus terreus* | Partial | 7 | 0 | 1 |
| PKI | 23617571 | *Emericella nidulans* | Partial | 7 | 0 | 1 |
| Pseurotin | 17722120 | *Aspergillus clavatus* | Perfect | 5 | 0 | 0 |
| Pseurotin | 17722120 | *Aspergillus fumigatus* | Partial | 5 | 17 | 0 |
| Pseurotin | 17722120 | *Aspergillus nomius* | Partial | 5 | 13 | 0 |
| Pseurotin | 17722120 | *Metarhizium anisopliae* | Partial | 5 | 11 | 0 |
| Pseurotin | 17722120 | *Metarhizium brunneum* | Partial | 5 | 9 | 0 |
| Pseurotin | 17722120 | *Metarhizium robertsii* | Partial | 5 | 10 | 0 |
| Pseurotin | 17722120 | *Penicillium arizonense* | Partial | 5 | 19 | 0 |

| | | | | | | |
|---|---|---|---|---|---|---|
| Pseurotin | 17722120 | *Penicillium solitum* | Perfect | 5 | 0 | 0 |
| Pyripyropene A | 21224862 | *Aspergillus fumigatus* | Partial | 8 | 0 | 1 |
| Pyripyropene A | 21224862 | *Neosartorya fischeri* | Partial | 8 | 0 | 1 |
| Radicicol | 18567690 | *Colletotrichum graminicola* | Perfect | 5 | 0 | 0 |
| Radicicol | 18567690 | *Colletotrichum sublineola* | Perfect | 5 | 0 | 0 |
| Roquefortine C | 22118684 | *Penicillium flavigenum* | Partial | 7 | 22 | 0 |
| Roquefortine C | 22118684 | *Penicillium rubens* | Not_found | | | |
| Roquefortine C | 22118684 | *Penicillium vulpinum* | Partial | 7 | 16 | 0 |
| TAN1612 | 21866960 | *Aspergillus kawachii* | Perfect | 5 | 0 | 0 |
| TAN1612 | 21866960 | *Aspergillus niger* | Perfect | 5 | 0 | 0 |

**Table S2: Prediction of secondary metabolism gene clusters with SMURF.**
Format as in table S1.

| Cluster name | Species name | SMURF | | | |
|---|---|---|---|---|---|
| | | **TAG** | **Size** | **Additional genes** | **Missing genes** |
| asperfuranone | *Aspergillus terreus* | Partial | 6 | 7 | 2 |
| asperfuranone | *Penicillium antarcticum* | Partial | 8 | 5 | 0 |
| aspernidineA | *Aspergillus calidoustus* | Partial | 6 | 1 | 0 |
| aspernidineA | *Emericella nidulans* | Partial | 6 | 16 | 0 |
| aurofusarin | *Fusarium poae* | Partial | 10 | 0 | 2 |
| aurofusarin | *Fusarium pseudograminearum* | Partial | 12 | 16 | 0 |
| bikaverin | *Fusarium fujikuroi* | Partial | 6 | 15 | 0 |
| bikaverin | *Fusarium mangiferae* | Partial | 6 | 12 | 0 |
| bikaverin | *Fusarium oxysporum* | Partial | 6 | 19 | 0 |
| bikaverin | *Fusarium verticillioides* | Partial | 6 | 13 | 0 |

| | | | | | |
|---|---|---|---|---|---|
| butenolide | *Fusarium avenaceum* | Not_found | | | |
| butenolide | *Fusarium graminearum* | Not_found | | | |
| butenolide | *Fusarium poae* | Not_found | | | |
| butenolide | *Fusarium pseudograminearum* | Not_found | | | |
| depudecin | *Colletotrichum orchidophilum* | Partial | 5 | 0 | 1 |
| depudecin | *Microsporum gypseum* | Partial | 5 | 0 | 1 |
| FDB2 | *Fusarium fujikuroi* | Not_found | | | |
| FDB2 | *Fusarium mangiferae* | Not_found | | | |
| FDB2 | *Fusarium proliferatum* | Not_found | | | |
| fumitremorgin | *Aspergillus fumigatus* | Partial | 8 | 25 | 1 |
| fumitremorgin | *Neosartorya fischeri* | Partial | 8 | 1 | 1 |
| fusaric_acid | *Fusarium fujikuroi* | Partial | 5 | 8 | 0 |
| fusaric_acid | *Fusarium mangiferae* | Partial | 5 | 14 | 0 |
| fusaric_acid | *Fusarium proliferatum* | Partial | 5 | 8 | 0 |
| fusarin | *Fusarium fujikuroi* | Partial | 9 | 13 | 0 |
| fusarin | *Fusarium proliferatum* | Partial | 8 | 9 | 1 |
| fusarubin | *Fusarium avenaceum* | Partial | 6 | 12 | 0 |
| fusarubin | *Fusarium fujikuroi* | Partial | 6 | 9 | 0 |
| fusarubin | *Fusarium mangiferae* | Perfect | 6 | 0 | 0 |
| fusarubin | *Fusarium oxysporum* | Partial | 6 | 5 | 0 |
| fusarubin | *Fusarium proliferatum* | Perfect | 6 | 0 | 0 |
| gibberellin | *Fusarium fujikuroi* | Not_found | | | |
| gibberellin | *Fusarium mangiferae* | Not_found | | | |
| gibberellin | *Fusarium proliferatum* | Not_found | | | |
| gliotoxin | *Aspergillus fumigatus* | Partial | 12 | 4 | 0 |

| | | | | | |
|---|---|---|---|---|---|
| gliotoxin | *Aspergillus udagawae* | Perfect | 12 | 0 | 0 |
| gliotoxin | *Neosartorya fischeri* | Partial | 12 | 12 | 0 |
| gliotoxin | *Neosartorya udagawae* | Perfect | 12 | 0 | 0 |
| HAS | *Arthroderma benhamiae* | Partial | 8 | 9 | 0 |
| HAS | *Arthroderma otae* | Partial | 8 | 5 | 0 |
| HAS | *Aspergillus fumigatus* | Partial | 8 | 10 | 0 |
| HAS | *Aspergillus udagawae* | Partial | 8 | 9 | 0 |
| HAS | *Neosartorya fischeri* | Partial | 8 | 10 | 0 |
| HAS | *Neosartorya udagawae* | Partial | 8 | 9 | 0 |
| HAS | *Trichophyton equinum* | Partial | 8 | 2 | 0 |
| HAS | *Trichophyton soudanense* | Partial | 8 | 4 | 0 |
| Helvonic acid | *Aspergillus fumigatus* | Not_found | | | |
| Helvonic acid | *Aspergillus udagawae* | Not_found | | | |
| Helvonic acid | *Metarhizium anisopliae* | Not_found | | | |
| Helvonic acid | *Metarhizium brunneum* | Not_found | | | |
| Helvonic acid | *Metarhizium guizhouense* | Not_found | | | |
| Helvonic acid | *Metarhizium robertsii* | Partial | 9 | 2 | 0 |
| Helvonic acid | *Neosartorya fischeri* | Not_found | | | |
| Neosartoricin | *Arthroderma benhamiae* | Partial | 5 | 14 | 0 |
| Neosartoricin | *Arthroderma otae* | Partial | 5 | 11 | 0 |
| Neosartoricin | *Aspergillus fumigatus* | Partial | 5 | 14 | 0 |
| Neosartoricin | *Microsporum gypseum* | Partial | 5 | 5 | 0 |
| Neosartoricin | *Neosartorya fischeri* | Perfect | 5 | 0 | 0 |
| Neosartoricin | *Trichophyton rubrum* | Partial | 5 | 25 | 0 |
| Neosartoricin | *Trichophyton soudanense* | Partial | 5 | 20 | 0 |

| | | | | | |
|---|---|---|---|---|---|
| Neosartoricin | *Trichophyton tonsurans* | Partial | 5 | 15 | 0 |
| Neosartoricin | *Trichophyton verrucosum* | Partial | 5 | 17 | 0 |
| Neosartoricin | *Trichophyton violaceum* | Partial | 5 | 12 | 0 |
| NG391 | *Metarhizium anisopliae* | Partial | 6 | 8 | 0 |
| NG391 | *Metarhizium brunneum* | Partial | 6 | 8 | 0 |
| NG391 | *Metarhizium guizhouense* | Partial | 6 | 11 | 0 |
| NG391 | *Metarhizium majus* | Partial | 6 | 9 | 0 |
| NG391 | *Metarhizium robertsii* | Partial | 6 | 9 | 0 |
| Patulin | *Aspergillus clavatus* | Partial | 13 | 10 | 2 |
| Patulin | *Penicillium expansum* | Perfect | 15 | 0 | 0 |
| Perithecial pigment | *Fusarium avenaceum* | Partial | 5 | 13 | 0 |
| Perithecial pigment | *Fusarium fujikuroi* | Partial | 5 | 10 | 0 |
| Perithecial pigment | *Fusarium mangiferae* | Partial | 5 | 1 | 0 |
| Perithecial pigment | *Fusarium oxysporum* | Partial | 5 | 6 | 0 |
| Perithecial pigment | *Fusarium proliferatum* | Partial | 5 | 1 | 0 |
| pigments | *Aspergillus clavatus* | Partial | 6 | 4 | 0 |
| pigments | *Neosartorya fischeri* | Partial | 6 | 6 | 0 |
| PKI | *Aspergillus rambellii* | Partial | 5 | 3 | 3 |
| PKI | *Aspergillus terreus* | Partial | 8 | 11 | 0 |
| PKI | *Emericella nidulans* | Partial | 7 | 0 | 1 |
| Pseurotin | *Aspergillus clavatus* | Not_found | | | |
| Pseurotin | *Aspergillus fumigatus* | Not_found | | | |
| Pseurotin | *Aspergillus nomius* | Partial | 5 | 12 | 0 |

| | | | | | |
|---|---|---|---|---|---|
| Pseurotin | *Metarhizium anisopliae* | Partial | 5 | 18 | 0 |
| Pseurotin | *Metarhizium brunneum* | Partial | 5 | 18 | 0 |
| Pseurotin | *Metarhizium robertsii* | Partial | 5 | 14 | 0 |
| Pseurotin | *Penicillium arizonense* | Partial | 5 | 22 | 0 |
| Pseurotin | *Penicillium solitum* | Partial | 4 | 11 | 1 |
| Pyripyropene A | *Aspergillus fumigatus* | Partial | 9 | 12 | 0 |
| Pyripyropene A | *Neosartorya fischeri* | Partial | 7 | 0 | 2 |
| Radicicol | *Colletotrichum graminicola* | Partial | 5 | 2 | 0 |
| Radicicol | *Colletotrichum sublineola* | Partial | 4 | 7 | 1 |
| Roquefortine C | *Penicillium flavigenum* | Perfect | 7 | 0 | 0 |
| Roquefortine C | *Penicillium rubens* | Perfect | 7 | 0 | 0 |
| Roquefortine C | *Penicillium vulpinum* | Partial | 7 | 7 | 0 |
| TAN1612 | *Aspergillus kawachii* | Partial | 5 | 2 | 0 |
| TAN1612 | *Aspergillus niger* | Partial | 5 | 1 | 0 |

**Table S3: Prediction of secondary metabolism gene clusters with ANTISMASH.** Format as in Table S1.

| Cluster name | Species name | ANTISMASH | | | |
|---|---|---|---|---|---|
| | | TAG | Size | Additional genes | Missing genes |
| asperfuranone | *Aspergillus terreus* | Partial | 8 | 12 | 0 |
| asperfuranone | *Penicillium antarcticum* | Partial | 8 | 10 | 0 |
| aspernidineA | *Aspergillus calidoustus* | Not_found | | | |
| aspernidineA | *Emericella nidulans* | Partial | 6 | 12 | 0 |
| aurofusarin | *Fusarium poae* | Partial | 12 | 4 | 0 |
| aurofusarin | *Fusarium pseudograminearum* | Partial | 12 | 27 | 0 |
| bikaverin | *Fusarium fujikuroi* | Partial | 6 | 15 | 0 |

| | | | | | |
|---|---|---|---|---|---|
| bikaverin | *Fusarium mangiferae* | Partial | 6 | 13 | 0 |
| bikaverin | *Fusarium oxysporum* | Partial | 6 | 12 | 0 |
| bikaverin | *Fusarium verticillioides* | Partial | 3 | 16 | 3 |
| butenolide | *Fusarium avenaceum* | Not_found | | | |
| butenolide | *Fusarium graminearum* | Not_found | | | |
| butenolide | *Fusarium poae* | Not_found | | | |
| butenolide | *Fusarium pseudograminearum* | Not_found | | | |
| depudecin | *Colletotrichum orchidophilum* | Partial | 6 | 6 | 0 |
| depudecin | *Microsporum gypseum* | Partial | 6 | 11 | 0 |
| FDB2 | *Fusarium fujikuroi* | Not_found | | | |
| FDB2 | *Fusarium mangiferae* | Not_found | | | |
| FDB2 | *Fusarium proliferatum* | Not_found | | | |
| fumitremorgin | *Aspergillus fumigatus* | Partial | 9 | 26 | 0 |
| fumitremorgin | *Neosartorya fischeri* | Partial | 9 | 14 | 0 |
| fusaric_acid | *Fusarium fujikuroi* | Partial | 5 | 10 | 0 |
| fusaric_acid | *Fusarium mangiferae* | Partial | 5 | 16 | 0 |
| fusaric_acid | *Fusarium proliferatum* | Partial | 5 | 10 | 0 |
| fusarin | *Fusarium fujikuroi* | Partial | 9 | 12 | 0 |
| fusarin | *Fusarium proliferatum* | Partial | 9 | 11 | 0 |
| fusarubin | *Fusarium avenaceum* | Partial | 6 | 12 | 0 |
| fusarubin | *Fusarium fujikuroi* | Partial | 6 | 13 | 0 |
| fusarubin | *Fusarium mangiferae* | Partial | 6 | 13 | 0 |
| fusarubin | *Fusarium oxysporum* | Partial | 6 | 15 | 0 |
| fusarubin | *Fusarium proliferatum* | Partial | 6 | 14 | 0 |
| gibberellin | *Fusarium fujikuroi* | Partial | 6 | 3 | 1 |

| | | | | | |
|---|---|---|---|---|---|
| gibberellin | *Fusarium mangiferae* | Partial | 6 | 3 | 1 |
| gibberellin | *Fusarium proliferatum* | Partial | 6 | 2 | 1 |
| gliotoxin | *Aspergillus fumigatus* | Partial | 12 | 12 | 0 |
| gliotoxin | *Aspergillus udagawae* | Partial | 12 | 5 | 0 |
| gliotoxin | *Neosartorya fischeri* | Partial | 12 | 15 | 0 |
| gliotoxin | *Neosartorya udagawae* | Partial | 12 | 3 | 0 |
| HAS | *Arthroderma benhamiae* | Partial | 7 | 6 | 1 |
| HAS | *Arthroderma otae* | Partial | 7 | 5 | 1 |
| HAS | *Aspergillus fumigatus* | Partial | 8 | 7 | 0 |
| HAS | *Aspergillus udagawae* | Partial | 7 | 2 | 1 |
| HAS | *Neosartorya fischeri* | Partial | 8 | 7 | 0 |
| HAS | *Neosartorya udagawae* | Partial | 8 | 6 | 0 |
| HAS | *Trichophyton equinum* | Partial | 7 | 4 | 1 |
| HAS | *Trichophyton soudanense* | Partial | 8 | 7 | 0 |
| Helvonic acid | *Aspergillus fumigatus* | Not_found | | | |
| Helvonic acid | *Aspergillus udagawae* | Partial | 7 | 4 | 2 |
| Helvonic acid | *Metarhizium anisopliae* | Partial | 9 | 17 | 0 |
| Helvonic acid | *Metarhizium brunneum* | Partial | 9 | 19 | 0 |
| Helvonic acid | *Metarhizium guizhouense* | Not_found | | | |
| Helvonic acid | *Metarhizium robertsii* | Partial | 9 | 20 | 0 |
| Helvonic acid | *Neosartorya fischeri* | Partial | 7 | 2 | 2 |
| Neosartoricin | *Arthroderma benhamiae* | Partial | 5 | 11 | 0 |
| Neosartoricin | *Arthroderma otae* | Partial | 5 | 14 | 0 |
| Neosartoricin | *Aspergillus fumigatus* | Partial | 5 | 9 | 0 |
| Neosartoricin | *Microsporum gypseum* | Partial | 5 | 11 | 0 |

| | | | | | |
|---|---|---|---|---|---|
| Neosartoricin | *Neosartorya fischeri* | Partial | 5 | 2 | 0 |
| Neosartoricin | *Trichophyton rubrum* | Partial | 5 | 25 | 0 |
| Neosartoricin | *Trichophyton soudanense* | Partial | 5 | 25 | 0 |
| Neosartoricin | *Trichophyton tonsurans* | Partial | 5 | 15 | 0 |
| Neosartoricin | *Trichophyton verrucosum* | Partial | 5 | 14 | 0 |
| Neosartoricin | *Trichophyton violaceum* | Partial | 5 | 11 | 0 |
| NG391 | *Metarhizium anisopliae* | Partial | 6 | 9 | 0 |
| NG391 | *Metarhizium brunneum* | Partial | 6 | 9 | 0 |
| NG391 | *Metarhizium guizhouense* | Partial | 6 | 11 | 0 |
| NG391 | *Metarhizium majus* | Partial | 6 | 10 | 0 |
| NG391 | *Metarhizium robertsii* | Partial | 6 | 9 | 0 |
| Patulin | *Aspergillus clavatus* | Partial | 14 | 2 | 1 |
| Patulin | *Penicillium expansum* | Partial | 8 | 7 | 7 |
| Perithecial pigment | *Fusarium avenaceum* | Partial | 5 | 13 | 0 |
| Perithecial pigment | *Fusarium fujikuroi* | Partial | 5 | 14 | 0 |
| Perithecial pigment | *Fusarium mangiferae* | Partial | 5 | 14 | 0 |
| Perithecial pigment | *Fusarium oxysporum* | Partial | 5 | 16 | 0 |
| Perithecial pigment | *Fusarium proliferatum* | Partial | 5 | 15 | 0 |
| pigments | *Aspergillus clavatus* | Partial | 6 | 12 | 0 |
| pigments | *Neosartorya fischeri* | Partial | 6 | 11 | 0 |
| PKI | *Aspergillus rambellii* | Not_found | | | |
| PKI | *Aspergillus terreus* | Partial | 7 | 8 | 1 |
| PKI | *Emericella nidulans* | Partial | 7 | 6 | 1 |

| | | | | | |
|---|---|---|---|---|---|
| Pseurotin | *Aspergillus clavatus* | Partial | 5 | 5 | 0 |
| Pseurotin | *Aspergillus fumigatus* | Partial | 5 | 19 | 0 |
| Pseurotin | *Aspergillus nomius* | Partial | 5 | 10 | 0 |
| Pseurotin | *Metarhizium anisopliae* | Partial | 5 | 19 | 0 |
| Pseurotin | *Metarhizium brunneum* | Partial | 5 | 12 | 0 |
| Pseurotin | *Metarhizium robertsii* | Partial | 5 | 20 | 0 |
| Pseurotin | *Penicillium arizonense* | Partial | 5 | 28 | 0 |
| Pseurotin | *Penicillium solitum* | Partial | 5 | 4 | 0 |
| Pyripyropene A | *Aspergillus fumigatus* | Partial | 9 | 11 | 0 |
| Pyripyropene A | *Neosartorya fischeri* | Partial | 9 | 4 | 0 |
| Radicicol | *Colletotrichum graminicola* | Perfect | 5 | 0 | 0 |
| Radicicol | *Colletotrichum sublineola* | Partial | 5 | 9 | 0 |
| Roquefortine C | *Penicillium flavigenum* | Partial | 7 | 8 | 0 |
| Roquefortine C | *Penicillium rubens* | Partial | 7 | 13 | 0 |
| Roquefortine C | *Penicillium vulpinum* | Partial | 7 | 9 | 0 |
| TAN1612 | *Aspergillus kawachii* | Partial | 5 | 11 | 0 |
| TAN1612 | *Aspergillus niger* | Partial | 5 | 15 | 0 |

**Table S4: List of 30 random fungal genomes used to compare Gecko3 and Evolclust.** The list include the species tag, species name and taxonomic classification.

| Species code | Species name | Taxonomy |
|---|---|---|
| ARTBE | Arthroderma benhamiae | Fungi;Ascomycota;Pezizomycotina;Eurotiomycetes;Onygenales;Arthrodermataceae;Arthroderma |
| ASPCR | Aspergillus cristatus | Fungi;Ascomycota;Pezizomycotina;Eurotiomycetes;Eurotiales;Aspergillaceae;Aspergillus |
| ASPFL | Aspergillus flavus | Fungi;Ascomycota;Pezizomycotina;Eurotiomycetes;Eurotiales;Aspergillaceae;Aspergillus |
| BIPOR | Bipolaris oryzae | Fungi;Ascomycota;Pezizomycotina;Dothideomycetes;Pleosporales;Pleosporaceae;Bipolaris |

| | | |
|---|---|---|
| CAPSE | Capronia semiimmersa | Fungi;Ascomycota;Pezizomycotina;Eurotiomycetes;Chaetothyriales;Herpotrichiellaceae;Phialophora |
| CERPL | Ceratocystis platani | Fungi;Ascomycota;Pezizomycotina;Sordariomycetes;Microascales;Ceratocystidaceae;Ceratocystis |
| COLTO | Colletotrichum tofieldiae | Fungi;Ascomycota;Pezizomycotina;Sordariomycetes;Glomerellales;Glomerellaceae;Colletotrichum |
| DICSQ | Dichomitus squalens | Fungi;Basidiomycota;Agaricomycotina;Agaricomycetes;Polyporales;Polyporaceae;Dichomitus |
| FIBRA | Fibroporia radiculosa | Fungi;Basidiomycota;Agaricomycotina;Agaricomycetes;Polyporales;Polyporaceae;Fibroporia |
| FUSPR | Fusarium proliferatum | Fungi;Ascomycota;Pezizomycotina;Sordariomycetes;Hypocreales;Nectriaceae;Fusarium |
| GIBMO | Fusarium verticillioides | Fungi;Ascomycota;Pezizomycotina;Sordariomycetes;Hypocreales;Nectriaceae;Fusarium |
| GRIFR | Grifola frondosa | Fungi;Basidiomycota;Agaricomycotina;Agaricomycetes;Agaricales;Schizophyllaceae;Grifola |
| HEBCY | Hebeloma cylindrosporum | Fungi;Basidiomycota;Agaricomycotina;Agaricomycetes;Agaricales;Cortinariaceae;Hebeloma |
| MELPE | Melanopsichium pennsylvanicum | Fungi;Basidiomycota;Ustilaginomycotina;Ustilaginomycetes;Ustilaginales;Ustilaginaceae;Melanopsichium |
| MICSP | Microsporidia sp. UGP3 | Fungi;Microsporidia |
| MICGY | Microsporum gypseum | Fungi;Ascomycota;Pezizomycotina;Eurotiomycetes;Onygenales;Arthrodermataceae;Microsporum |
| MUCCI | Mucor circinelloides | Fungi;Mucoromycota;Mucoromycotina;Mucoromycotina_C;Mucorales;Mucoraceae;Mucor |
| PENAR | Penicillium arizonense | Fungi;Ascomycota;Pezizomycotina;Eurotiomycetes;Eurotiales;Aspergillaceae;Penicillium |
| PENCO | Penicillium coprophilum | Fungi;Ascomycota;Pezizomycotina;Eurotiomycetes;Eurotiales;Aspergillaceae;Penicillium |
| PENNO | Penicillium nordicum | Fungi;Ascomycota;Pezizomycotina;Eurotiomycetes;Eurotiales;Aspergillaceae;Penicillium |
| PENST | Penicillium steckii | Fungi;Ascomycota;Pezizomycotina;Eurotiomycetes;Eurotiales;Aspergillaceae;Penicillium |
| PNEJI | Pneumocystis jirovecii | Fungi;Ascomycota;Taphrinomycotina;Pneumocystidomycetes;Pneumocystidales;Pneumocystidaceae;Pneumocystis |
| PSEBR | Pseudozyma brasiliensis | Fungi;Basidiomycota;Ustilaginomycotina;Ustilaginomycetes;Ustilaginales;Ustilaginaceae;Pseudozyma |
| PYRTR | Pyrenophora tritici-repentis | Fungi;Ascomycota;Pezizomycotina;Dothideomycetes;Pleosporales;Pleosporaceae;Pyrenophora |
| RHYAG | Rhynchosporium agropyri | Fungi;Ascomycota;Pezizomycotina;Leotiomycetes;Helotiales;Helotiales_F;Rhynchosporium |
| YEAST | Saccharomyces cerevisiae | Fungi;Ascomycota;Saccharomycotina;Saccharomycetes;Saccharomycetales;Saccharomycetaceae;Saccharomyces |

| | | |
|---|---|---|
| SCEAP | Scedosporium apiospermum | Fungi;Ascomycota;Pezizomycotina;Sordariomycetes;Microascales;Microascaceae;Scedosporium |
| SPOSC | Sporothrix schenckii | Fungi;Ascomycota;Pezizomycotina;Sordariomycetes;Ophiostomatales;Ophiostomataceae;Sporothrix |
| TALCE | Talaromyces cellulolyticus | Fungi;Ascomycota;Pezizomycotina;Eurotiomycetes;Eurotiales;Trichocomaceae;Talaromyces |
| ZYMTR | Zymoseptoria tritici IPO323 | Fungi;Ascomycota;Pezizomycotina;Dothideomycetes;Capnodiales;Mycosphaerellaceae;Zymoseptoria |

**Table S5: Comparison between clusters predicted in Evolclust and in different runs of Gecko3.**

Columns indicate results for different runs of Gecko3. Rows indicate number of clusters predicted by both methods, number of identical clusters and number of cluster pairs that overlap in at least a given percentage.

| Shared clusters type | Evolclust vs gecko-q3d3 | Evolclust vs gecko-q4d3 | Evolclust vs gecko-q5d3 | Evolclust vs gecko-q4d5 |
|---|---|---|---|---|
| Total clusters Evolclust | 6482 | 6482 | 6482 | 6482 |
| Total clusters gecko | 7468 | 6603 | 5528 | 7184 |
| All shared | 5341 | 5011 | 4299 | 5227 |
| Identical | 865 | 775 | 630 | 848 |
| 100% | 3349 | 3183 | 2740 | 3126 |
| 90% | 110 | 94 | 75 | 161 |
| 75% | 433 | 419 | 370 | 498 |
| 50% | 381 | 357 | 339 | 377 |
| 30% | 203 | 183 | 145 | 217 |
